# Filter-banks and artificial intelligence in seizure detection using electroencephalograms

**DOI:** 10.1101/105650

**Authors:** M. A. Pinto-Orellana, F. R. Cerqueira

## Abstract

Epilepsy is the most typical neurological disease in the world, and it implies an expensive and specialized diagnosis process based on electroencephalograms and video recordings. We developed a method that only requires the brainwave provided by the difference between two standard-located electrodes. Our proposed technique separates the original signal using a filter array with three different types of filters, and then extracts several features based on information theory and statistical information. In our study, we found that only 10 characteristics, of which the most important are related to higher frequencies, are required to offer an accuracy of 94%, a specificity of 95% and a sensitivity of 87% using C4.5 decision trees.

## Introduction

Epilepsy is one of the most common and critical neurological diseases. The World Health Organization estimates that between 50 and 200 million people worldwide has some degree of epilepsy [1]. The diagnosis and treatment of this illness used to be an expensive and long process [2]. Consequently, in order to reduce the time, cost, and specialist-dependency of a diagnosis various alternatives were proposed [3].

The typical diagnostic method is to analyze the brainwaves or electroencephalograms (EEGs) and use video recordings to determine the actual patient state [4]. Several researches in the field shown that it is possible to detect seizures using only the brainwaves recorded in the EEGs using different techniques that range from analog circuitry [5] to artificial intelligence approaches [6].

EEGs could be defined as electrical signals measured in standard points of the brain at the scalp. In normal conditions, EEG values oscillate between 5 and 50 microvolts in a bandwidth frequency from 0 to 100 Hz [7]. There are two main issues with brainwaves signal processing: Noise and subject data dependency. The lower values of electrical tension of the EEGs made them vulnerable to interference originated in the equipment or, even by the patient itself. The EEGs are so sensitive that even a muscular movement could generate some vibrations at the scalp that could lead to misinterpret the brain electrical signal [8, 9].

The second usual issue with EEGs is their variations from one patient to another one. Some techniques of normalization of EEG are commonly extended, like statistical normalization or spectral normalization. But, according to the objective of the processing of the brainwaves, these methods could reduce the efficiency of the proposed techniques [10]. Because of that, several algorithms are prepared to work only with one patient at the same time. This leads to methods that require that the patient need to personally train the algorithm with its own data before be able to use it.

Our proposed method relies in two well-known areas of knowledge: Digital signal processing, and artificial intelligence. Initially, we describe the basis of our system: The techniques to obtain the features of the original time series, the feature selection, and the machine learning algorithm that we used. And, in a second part, we described in depth the features that we found that are enough to recognize efficiently the seizure instants.

## Data source

To obtain EEGs from patients who suffered epilepsy is not an easy task. Thus, we relied in a publicly available and trustable large dataset: the Children Hospital Boston – Massachusetts Institute of Technology (CHBMIT) dataset from the Physionet project [11, 12]. This database stores recordings from 23 different child patients from an average duration of 24 hours for each one. Each recording was made following the 10-20 standard layout system for EEG and each one was sampled with a frequency of 256Hz.

For this research, we only use the signal from the electrode FP7 (left front-parietal area) using the reference of the electrode of the position P7 (left parietal lobe).

## Method

We separated our system in three subsystems: frequency splitting, feature extraction and analysis, and the detection subsystem by itself.

**Fig. 1.**
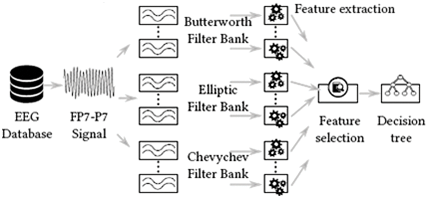
Workflow of the proposed method.

### A. Frequency splitting subsystem

Due to the EEGs save their information in the form of frequency variations, it is common that time-frequency approaches are used to process this kind of signals. Low-pass filtering data is a classic procedure before signal processing because this removes the high frequency noise. Nevertheless, the quality of the output signal depends on the type of the filter and its order.

Here, we applied a different methodology: We used a filter bank, an array of filters for different frequency intervals, like the idea behind wavelets transform (WT) [13]. However, in our case we maintain the filtered signal rather than get seed codes as WT does. Our proposal is to separate the complete signal in ten segments with a frequency bandwidth of 12.5Hz each one. Moreover, there are several types of filters that can be applied with different performance results in frequency and phase response. Thus, we selected three different second-order filtering methods: Butterworth (BTW), Chevychev (CHY), and elliptic (ELP) filters.

### B. Feature analysis subsystem

Thus far, we separated one signal into 30 time series. Later, the next phase is to obtain features of these series to be processed. One typical approach is to regrouping the data in small intervals to obtain several characteristics. In our method, we use a window of one second, or 256 samples, to calculate the features.

Two types of features were selected: Statistical and theoretical-information characteristics. For each data slide, we calculated 17 features that are detailed in Table 1. Hence, we obtained 510 features in total per second. They are a huge quantity of features, but also computational time feasible for a learning machine algorithm.

**Table 1:**
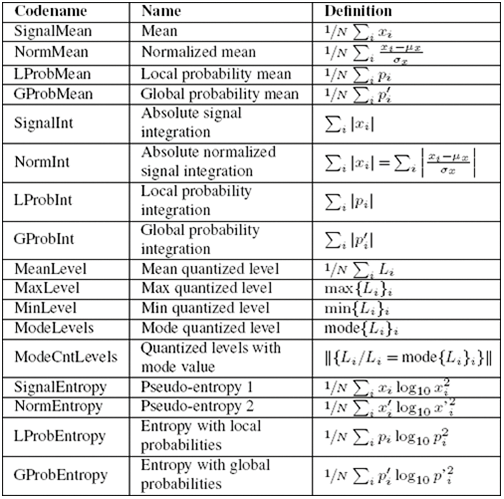
List of the features. It should be noted that *N* is the number of samples in the window, *x*_*i*_ is a sampled point, 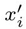 is a normalized sampled point (according to the estimated mean *μ*_*x*_ and the standard variation *σ*_*x*_), *p*_*x*_ is the probability of a value to appear in the window, and 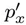 is the estimated probability of a value to appear in the whole signal.

### C. Detection subsystem

There are several machine learning algorithms that perform very well with EEG data, like random forests, and support vector machines. But they lack an easy interpretation of the data and act like black-box algorithms. In our study, we prefer to use a C4.5 decision tree as a base algorithm for two reasons: First, it could be easily translated into small programs based on conditionals, and the second, and foremost, reason it can offer a human-understandable interpretation of the data.

We have 510 features per instance, but we know that for several kinds of datasets, it is usual to use non-linear transformations of the data for improving the accuracy of the algorithm during the training. Thus, we used for transformations that are shown in Table 2. After that, our dataset has 2040 features per instance, hence we tried to reduce this number in order to find the most representatives and offer a better understanding of the process.

**Table 2:** Transformations of the signal.

| Transformation | Definition |
| --- | --- |
| Linear | $f(x) = x$ |
| Logarithmic | $f(x) = \log_{10} x^2$ |
| Exponential | $f(x) = \exp(x^{0.5})$ |
| Tangential hyperbolic | $f(x) = \tanh(x^{0.5})$ |

To find the minimum quantity of characteristics, we randomly split ten times the dataset in a proportion of 60% for training, and 40% for testing. Later, we use our classification algorithm as a predictor of the efficiency of each column of the data set, assigning the accuracy and recall of each feature as its score, and after sorting the columns according that value, we greedily add each column until the overall performance was not improved. Using this procedure, we found the ten most representative columns.

## Results

As it was mentioned before, we compiled the signals of the 24 subjects that are included in the CHBMIT dataset. It should be noted that our system relies in the libraries and environments provided by the Weka software (version 3.7.12) and the Numpy 1.11.1, Scipy 0.17.1, and Scikit-learn 0.17.1 libraries.

To validate our proposal, we split the whole data in ten sets of training and test subsets randomly selected with a proportion of 60% and 40%, respectively. In this way, we ensure the response of the system with an average training of approximately 7058.08 seconds of seizures, and a testing with nearly 4706.36 seconds with seizure events. After performing the feature selection, we find 10 representative characteristics that are shown in the Table 3. And, overall in our testing environment, we obtained an accuracy close to 94%, and a true positive rate near to 87%, with a specificity of 95%.

**Table 3:** Most representative features.

| Bandwidth frequency [Hz] | Filter type | Feature Linear | Transformation | Relevance |
| --- | --- | --- | --- | --- |
| 12.5–25.0 | BTW | LProbInt | Linear | 6th |
| 25.0–37.5 | BTW | LProbInt | Linear | 4th |
|  | BTW | LProbMean | Linear | 7th |
| 37.5–50.0 | BTW | ModeLevels | Linear | 2nd |
|  | BTW | GProbEntropy | Linear | 8th |
|  | BTW | NormMean | Linear | 10th |
| 75.0–87.5 | BTW | MaxLevels | Linear | 9th |
| 87.5–100.0 | ELP | LProbMean | Logarithmic | 3rd |
|  | BTW | SignalEntropy | Linear | 5th |
| 112.5–125.0 | CHY | GProbEntropy | Exponential | 1st |

## Discussion

The literature of EEG commonly associates the frequency bands lower than 50Hz to several conditions or activities, whilst the waves with higher frequency are not clearly associated with any particular condition. In our results, we found a surprising fact: The most important characteristic needed a non-linear transformation, and above all, it was related to the highest analyzed band. Also, from the best 10 features, 4 (40%) were in bands higher than 50Hz, normally ignored in another kind of EEG processing systems. But, we have no enough information to link any abnormal behavior in these higher frequencies with seizures.

Regarding to the type of filters, we noted that a second-order filter was sufficient to provide the vast majority of the relevant features of the system. However, the three most important characteristics were obtained from the three kinds of filters proposed. Thus, we cannot ignore them in the overall analysis.

## Conclusion

We proposed a new kind of seizure detection system with uses time series features obtained from filter banks. Our proposed method demonstrates four main benefits: First, it uses only two electrodes that allow a practical physical implementation. Second, it has a good performance, reaching an accuracy greater than 94%, a sensitivity close to 87%, and a specificity near to 95%. Third, using decision trees we are able to translate the prediction tree in programs that can be stored in small devices. And, fourth, our system was designed to work with an independence of the origin of the data, that is, we could generalize the system to work with different patients being previously trained with other ones.

## Acknowledgment

This work has been supported by grants from Fundação de Amparo à Pesquisa de Minas Gerais (FAPEMIG) and the PAEC agreement between the Organization of American States (OAS) and Universidade Federal de Viçosa (UFV).

Marco A. Pinto-Orellana and Fabio R. Cerqueira (*Departamento de Informática, Universidade Federal de Viçosa, Brazil*)

